# Population studies of the wild tomato species *Solanum chilense* reveal geographically structured major gene-mediated pathogen resistance

**DOI:** 10.1101/2020.05.29.122960

**Authors:** Parvinderdeep S. Kahlon, Shallet Mindih Seta, Gesche Zander, Daniela Scheikl, Ralph Hückelhoven, Matthieu H. A. J. Joosten, Remco Stam

**Author notes:** Author for correspondence Remco Stam, Chair of Phytopathology, TUM School of Life Sciences, Technical University of Munich, Emil-Ramann-Str. 2, 85354, Freising, Germany. Funding: This work was funded through the German Science Foundation (DFG), SFB924.

## Abstract

Natural plant populations encounter strong pathogen pressure and defense-associated genes are known to be under different selection pressure dependent on the pressure by the pathogens. Here we use wild tomato *Solanum chilense* populations to investigate natural resistance against *Cladosporium fulvum*, a well-known pathogenic fungus of domesticated tomatoes. We show that populations of *S. chilense* differ in resistance against the pathogen. Next, we explored the underlying molecular processes in a species wide-context. Then, focusing on recognition of the two prominent avirulence factors secreted by *C. fulvum* (Avr4 and Avr9) in central and northern populations of *S. chilense* we observed high complexity in the cognate *homologues of Cladosporium resistance* (*Hcr9*) locus underlying the recognition of these effectors. Presence of canonical genomic regions coding for *Cf-4* and *Cf-9*, two major dominant resistance genes in the *Hcr9* locus recognizing Avr4 and Avr9, respectively, does not meet prediction from Avr response phenotypes. We find both genes in varying fractions of the plant populations and we show possible co-existence of two functionally active resistance genes, previously thought to be allelic. Additionally, we observed the complete local absence of recognition of additional Avr proteins of *C. fulvum*. In the southern populations we attribute this to changes in the coregulatory network. As a result of loss of pathogen pressure or adaptation to extreme climatic conditions. This may ultimately explain the observed pathogen susceptibility in the southern populations. This work puts major gene mediated disease resistance in an ecological context.

## Introduction

How plants and their pathogens adapt to one another in their natural habitat is still poorly understood. Some studies highlight the local adaptations in context to plant-pathogen interactions: the wild flax-flax rust pathosystem is such an example, where more resistant wild flax harbored more virulent strains of the rust (Thrall *et al.*, 2002; Thrall & Burdon, 2003). Similar mechanisms of the co-occurrence of virulent strains of the powdery mildew *Podosphaera plantaginis* and more resistant plants of *Plantago lanceolata* have also been documented (Laine, 2005; Soubeyrand *et al*., 2009). Furthermore, complex multi-host and multi-pathogen systems, with clear differences at a regional scale, have been observed in anther smut- fungi-infecting *Dianthus* hosts in the southern European Alps (Petit *et al*., 2017). The molecular mechanisms that is thought to drive these interactions is a gene-for-gene interaction (Flor, 1971), where single gene encoded pathogen molecules, now referred to as effectors, are recognized by specific receptors present in resistant host plants, also known as major resistance genes. The need to recognize an invading pathogen and subsequently evade the recognition by the plant and pathogen, respectively, leads to a constant co-evolutionary arms race, often described as the Red Queen dynamics (Van Valen, 1973). These dynamics are thought to have led to the plethora of different defense related genes in plants as we know it today. Wild crop relatives have been used as a gene pool for isolating and introducing genetic resistance against many different fungal pathogens. Genetic diversity of natural populations against microbes have been explored by sequence analysis or by experimental biology often still as independent, though complementary, approaches (Salvaudon *et al*., 2008), leading to a huge gap in our understanding of local adaptations and their role in shaping the current diversity of such resistance genes in an ecological context.

Tomato leaf mold is caused by the non-obligate biotrophic fungus *Cladosporium fulvum* syn. *Passalora fulva*. At the time of infection, *C. fulvum* enters through stomata into the leaf and colonizes the apoplastic spaces (Stergiopoulos & de Wit, 2009). The fungus secretes various small proteins, also referred to as effectors or avirulence factors (Avrs) when being recognized by a resistant host, into the apoplast with the aim to manipulate the host for its successful colonization of the host tissue. The infection leads to severe yellowing and wilting of the leaves, which ultimately leads to a loss of photosynthetic capacity and thus a loss in yield or reduced reproducible fitness. *C. fulvum* is a globally occurring pathogen with a clear genetic diversity (Iida *et al*., 2015). It is thought to originate in the Andean region, where it likely has co-evolved with one or several of wild tomato species that inhabit a mountain range from central Ecuador to northern Peru (de Wit *et al*., 2012). During evolution, as a result of selection pressure imposed by virulent strains of *C. fulvum*, several of the wild tomato species have evolved resistance genes whose products mediate recognition of the Avrs secreted by *C. fulvum* (Joosten & de Wit, 1999). This recognition facilitates host resistance following the gene-for-gene model. Resistance is eventually achieved when Avr-activated defense leads to a hypersensitive response (HR), which includes programmed cell death (PCD). This localized PCD is associated with various additional defense responses such as a massive callose deposition, and prevents *C. fulvum* from obtaining nutrients from the host, thereby limiting further pathogen ingress and multiplication (Lazarovits & Higgins, 1976).

To date, 24 *C. fulvum* (*Cf*) resistance genes have been mapped (Kanwar *et al*., 1980). Due to the complexity of the *Cf* genes and their loci, only a small number of *Cf* genes have been cloned and verified for their functionality. *Cf* genes are highly repetitive in their leucine-rich repeat encoding parts and functional homologs are often accompanied by non-functional ones, with very little sequence variation between them (Kruijt 2005). *Cf-9* from *S. pimpinellifolium* encodes a cell surface receptor-like protein (RLP) and was the first *Cf* resistance gene to be cloned (Jones *et al*., 1994). The *Cf-9* gene product recognizes the Avr9 protein of *C. fulvum*, which is a highly stable, cysteine-knotted peptide of unknown function (Scholtens-Toma & de Wit 1988). The *Cf-9* gene belongs to the *Hcr9* (homologs of *C. fulvum* resistance gene *Cf-9*) gene cluster, which is located on chromosome 1. In addition to *Cf-9, Cf-4* is another well-studied *Cf* gene from the *Hcr9* cluster. The *Cf-4* gene product recognizes Avr4 of *C. fulvum*, which is a chitin-binding protein having eight cysteine residues (Joosten et al., 1994; van den Burg et al., 2004), and the *Cf-4* gene originated from *S. habrochaites* (Thomas *et al*., 1997). Studies on MM-Cf9 which is an Avr9- recognising introgression line of the domesticated tomato, *S. lycopersicum* cv Money Maker, revealed presence of five *Cf-9* homologs, *Hcr9*-*9A* to *Hcr9*-*9E*, mapped at the short arm of chromosome 1, with *Hcr9-9C* being the functional *Cf-9* gene (Parniske *et al*., 1997). Similarly, *Hcr9*-*4A* to *Hcr9*-*4E* is present in the Avr4-recognising MM-Cf4 and *Hcr9*-*4D* corresponds to *Cf-4* (Thomas *et al*., 1997). *Cf-4* and *Cf-9* lie at the same locus on chromosome 1 and are assumed to be mutually exclusive or even allelic. Crossings between recombinant inbred lines carrying *Cf-4* and *Cf-9* resulted in extreme genetic instability in the offspring (Parniske *et al.*, 1997, Thomas *et al.*, 1997).

Appreciating the important roles of these *Cf* genes and their assumed role in co-evolution between wild tomato and native *C. fulvum*, it is surprising that only a few studies have sought to investigate *Cf* gene diversity. An effectoromics approach was exploited by Mesarich *et al.* (2017), to identify plants carrying novel *Cf* genes to be potentially used in plant breeding programs in the future. Studies have shown that in *S. pimpinellifolium* several putative homologs are present, but their function remains unknown (Caicedo *et al*., 2004; Caicedo, 2008). Another study identified four variants of *Cf-9* (originally isolated from *S. pimpinellifolium*), from its close relative *S. habrochaites* and each one variant of *Cf-4* from *S. habrochaites, S. chilense, S. chmielewskii, S. neorickii* and *S. arcanum* (Kruijt *et al*. 2005). In-spite of multiple single nucleotide polymorphisms (SNPs) being present in the isolated *Cf* gene variants, all of them showed the ability to induce an HR after recognition of Avr9 and Avr4, respectively, which led the authors to conclude that Avr4 and Avr9 recognition is conserved among wild tomato. However, a species-wide analysis of different accessions of *S. pimpinellifolium* revealed intragenic recombination to have occurred between *Hcr9-9D* and *Hcr9-9C*/*Cf-9.* The resulting allele, *9DC*, does not co-exist with the original *Cf-9* allele in the individual plants. Variant *9DC* is the more common allele in the species and the product also recognizes Avr9 (Van der Hoorn *et al*., 2001), indicating that one likely cannot speak of conservation of *Cf* alleles *sensu stricto.* In addition, plants that recognize both Avr4 and Avr9 have not been identified. Detailed knowledge on the relationship between *Cf-4* and *Cf-9* in other accessions or wild populations of tomato, and on their actual roles in resistance is still lacking.

To perform more detailed studies on resistance provided by *Cf* genes in an ecological context, we selected *S. chilense*, one of seventeen wild tomato species, as it covers a wide variety of habitats on the western slopes of the Andes, ranging from Peru to Northern Chile (Nakazato *et al*., 2010). The species range spreads from the edges of the Atacama Desert, as the southern edge of the range, to relatively wet, high altitude regions (up to 3500 meter above sea level), as well as in very specific coastal regions that experience regular sea fog (resulting in a relatively high humidity) in the most northern part of Chile, as well as in south Peru (Cereceda & Schemenauer 1991; Peralta *et al*., 2008; Chetelat *et al*., 2009). This highly varied distribution results in sub- populations of this species that encounter different environmental challenges. Ultimately, these different habitats lead to genetic differentiation within the species. The *S. chilense* population can be clustered in four groups: north, central, southern highlands and southern lowlands. The southern highlands and southern lowlands populations are derived from the central group (Böndel*et al*., 2015). This divergence was confirmed by whole genome sequencing, and multiple sequential markovian coalescent simulations revealed that migrations happened from the central group southward 50,000 to 200,000 years ago (Stam *et al*., 2019b). The strong differentiation between habitats leads to clearly observable adaptations. Southern populations respond faster to drought (Fischer *et al*., 2013) whereas high altitude populations are more cold-tolerant (Nosenko *et al*., 2016). In addition to adaptations to abiotic factors, these habitats are expected to be home to different biotic stressors at different intensity levels, including various pathogen species, which is anticipated to result in genetic variation in pathogen defense-associated genes. Indeed, differences in resistance properties of the various *S. chilense* populations against three pathogens, *Alternaria solani, Phytophthora infestans* and a *Fusarium* spp, are observed (Stam *et al*., 2017). Moreover, large genetic variation has been observed within *S. chilense* populations in another resistance-associated gene family; the nucleotide-binding leucine-rich repeat (NLR) resistance genes. These genes show clear presence-absence-variation when compared to related tomato species (Stam *et al*., 2019a), and an in-depth resequencing study, covering the whole species range of *S. chilense*, shows evidence that the selection pressure imposed on the individual NLRs differs for each of the populations (Stam *et al*., 2019b).

Little data exists showing biological interactions between *C. fulvum* and wild tomato species. In this study we investigated the interaction of the fungal pathogen *C. fulvum* with the wild tomato species *S. chilense* throughout the geographical range of the species. We show that *S. chilense* plants from different locations first of all show differences in their resistance to the fungus, including complete loss of resistance in some populations. By investigating the well characterized genes of the *Hcr9* locus, we furthermore place major gene mediated immunity in an ecological context.

## Materials and methods

### Plants and fungal material

Seeds of the fifteen accessions (populations) of *S. chilense* used in our studies were originally obtained from the C. M. Rick Tomato Genetics Resource Center of the University of California, Davis (TGRC UC-Davis) (http://tgrc.ucdavis.edu/). The selected accessions were LA1958, LA1963, LA2747, LA2931, LA2932, LA2750, LA2959, LA3111, LA3784, LA3786, LA4107, LA4117A, LA4118 and LA4330. Each Accession number represents a random collection of seeds from a wild population and is propagated by TGRC in a way to maintain maximum genetic diversity. Böndel et al (2015) have shown that this seed multiplication has negligible effect on the genetic diversity of the accessions. Hence, each accession can be considered to truly represent the diversity in the wild plant populations. Introgression lines *S. lycopersicum* cv Moneymaker (MM) of *Cf-9, Cf-4* and *Cf-5* were generated by Tigchelaar *et al.* (1984) and *S. pimpinellifolium* (LP12), in which the *9DC* was identified, and which was used as a control in all our assays were provided by TGRC UC-Davis. Per *S. chilense* accession 8-17 plants were grown. Plants were grown in a controlled glasshouse, with a minimum daytime temperature of 24°C and 16 hours of light and 8 hours of dark conditions. Adult plants were cut back at bi- weekly intervals to assure the presence of mature, fully developed, yet not senescent, leaves during each repetition of the experiment.

A race 5 strain of *C. fulvum* was maintained on 1/2 potato dextrose agar (PDA) medium and incubated at 16°C in the dark.

### Visualization of infection phenotype and quantification of *C. fulvum* biomass from tomato leaves

Spray inoculation of *C. fulvum* at a concentration 20,000 conidia/ml was performed on 3-week old plant cuttings of population LA3111, LA4330, MM-Cf-9 and MM-Cf-5 plants; water-inoculated plants served as a negative control in the experiment. Plants were maintained at 24°C and 16 hours of light and 8 hours of dark, with 100% humidity for the first two days, after which 80% humidity was maintained throughout the experiment.

Photographs of leaflets of inoculated plants were collected at 14 days post inoculation (dpi) and Microscopy was performed from 7dpi to 19dpi on bleached leaves following staining in acetic acid (25%) 1:9 + ink (K□nigsblau, Pelikan, 4001) then washed in water and analysed under brightfield microscope (Zeiss, Imager Z1m).

For quantification, leaflets of inoculated plants were collected at 14 days post inoculation (dpi) and DNA was isolated using the protocol published by Yan *et al*. (2008). The infection loads or approximate amounts of *C. fulvum* DNA present in inoculated leaves were quantified by qPCR using the DNA-binding fluorophore Maxima SYBR Green Master Mix (2X) with ROX solution (Thermo Scientific). All qPCR reactions were performed on an AriaMx Real-Time PCR system (Agilent Technologies, Waldbronn, Germany). PCR were performed with the primer pairs RS158 (5’-GTCTCCGGCTGAGCAGTT-3’)/ ITS4 (5’-TCCTCCGCTTATTGATATGC-3’) and RS001 (5’-GCCTACCATGAGCAGCTTTC-3’)/ RS002 (5’-CAATGCGTGAGAAGACCTCA- 3’), annealing to the ITS region and *elongation factor 1 alpha*, respectively, present in the *C. fulvum* DNA. Templates used for this experiment were genomic DNA isolated from the leaves inoculated with the pathogen and water. Amplifications using both primer pairs for each sample were done on the same plate and in the run. The experiment was performed on three plants of MM-Cf-9 and MM-Cf-5 each and five plants of LA3111 and LA4330 each. All the plants samples well evaluated in three technical replicates and Cq differences higher than one among technical replicates was not considered in the final evaluation. The Cq values and the linear equations from the sensitivity graphs were used to calculate the quantities of pathogen and plant DNA in each sample (Figure S1).

### Apoplastic washing fluid (AF), Avr9 and Avr4 infiltration assay

AF containing the complete set of *C. fulvum* Avrs, except for Avr5, was obtained by isolating AF from leaves of MM-Cf-5 plants colonized by race 5 of *C. fulvum*, at 10 to 14 days after inoculation. Furthermore, a preparation of Avr9 concentrated from AF by acetone precipitation, leaving the Avr9 peptide in the supernatant, and Avr4 produced in the yeast *Pichia pastoris*, were employed.

Using a 1 ml syringe (Braun, Omnifix) without a needle, the AF, Avr9 and Avr4 preparations were infiltrated from the lower side of fully expanded leaves of the different populations. Experiments were performed in three independent biological replicates. Infiltrations were done in fully expanded leaves of the same adult plant, with two technical replicates per plant. Readings were performed between 2 to7 days post infiltration. For Avr9 infiltration, an HR was typically observed within 1-4 days, whereas the AF-and Avr4-triggered HR were observed later. Readings were not performed later than 7 days after infiltration.

### Presence of *Cf-9*-, *9DC*-and *Cf-4*-specific regions in different *S. chilense* populations

Screening of *S. chilense* populations for the presence of gene-specific regions of *Cf-9, 9DC* and *Cf-4* was performed through PCR amplification using the gene-specific primers CS5-CS1 and DS1-CS1 for *Cf-9* and *9DC*, respectively (Van der Hoorn *et al*., 2001). Primer pair PSK047 (5’- ACGACAGAAGAACTC-3’)/ PSK050 (5’-GATGGAATTGGTCCTT-3’), was designed to amplify the canonical *Cf-4* domain (Fig. S2). DNA was isolated using standard CTAB extraction, using the same samples as used in a previous study (Stam *et al.*, 2016; Stam *et al.*, 2019b). PCR on gDNA was performed using Promega Green GoTaq® Polymerase and the products were analyzed by 1% agarose gel electrophoresis. As a PCR control *elongation factor 1 alpha*, amplified with the primer pair 5’-GTCCCCATCTCTGGTTTTGA-3’/ 5’- GGGTCATCTTTGGAGTTGGA-3’, was included and MM-Cf-9 and MM-Cf-4, and LP12 served as positive and negative controls, respectively.

### Semi-quantitative evaluation of the expression of a *Cf-9* homolog in the southern population LA4330

We evaluated the transcript levels of the *Cf-9* homolog in LA4330 at eight hours after Avr9 or water infiltration. RNA from the infiltrated leaves was extracted using the RNeasy plant mini kit (Qiagen) and cDNA was synthesized using the QuantiTect reverse transcription kit (Qiagen). Amplification of *Cf-9* the canonical region of the *Cf-9* homolog was performed with primer pair binding on the start and end of the ORF, PSK009 (5’-ATGGATTGTGTAAAACTTGTATTCCT -3’) /PSK010 (5’-CTAATATCTTTTCTTGTGCTTTTTCA -3’), and the product was visualized on 1% agarose gel.

### Whole genome sequence analyses for *Cf* co-receptors in *S. chilense*

We extracted the sequences for four important components of the Cf protein signaling complex, for which we chose SUPPRESSOR OF BIR1 (SORBIR1), SOMATIC EMBRYOGENESIS RECEPTOR-KINASE 3A (SERK3a) and AVR9/CF-9-INDUCED KINASE 1 (ACIK1) (Wu, 2020), as well as for an additional potential co-receptor of the BAK1 signaling complex, BIR2 (Halter *et al*., 2014) from NCBI. To identify the genomic sequences encoding these co-receptors and regulators of Cf proteins in *S. chilense*, we performed a BLAST search against the *S. chilense* reference genome (Stam *et al*., 2019b). Each *SOBIR1, SEKR3a* and *ACIK1* yielded one unequivocal best target sequence in the reference genome and visual inspection after alignment confirmed that the *S. chilense* homologs are in all cases similar to the reference gene of *S. lycopersicum. BIR2* yielded two good hits, of which we dubbed the second one *BIR2b*. In order to look for polymorphisms in three plants, one from LA3111, and one each from southern populations LA2932 and LA4330 (Stam, *et al*. 2019a), we obtained the genomic coordinates of the genes of interest from our reference genome and called the SNPs in the region of interest using samtools mpileup and bcftools call (-mv -Oz) (Li *et al*., 2009; Li, 2011). We removed low quality indels and the resulting vcf files were tabulated using tabix, to allow the consensus sequences for the genomic regions to be extracted using bcftools consensus. Finally, we used gffread from gffutils (Pertea and Pertea 2020) to obtain the correct coding sequence for each of the plants. All multiple sequence alignments were made and inspected using aliview (Larsson, 2014).

### Statistical testing

ANOVA, with post hoc Tukey honest significant difference test was performed using R command Tukey HSD (anova) to test the statistical significance in pathogen load detected in the inoculated plants via qPCR.

## Results

### *S. chilense* populations show differences in resistance against *C. fulvum*

To confirm that *C. fulvum* is capable of successfully infecting *S. chilense* and to test whether *S. chilense* populations show differences in their resistance spectrum to *C. fulvum*, we spray-inoculated randomly selected individual plants of *S. chilense* from geographically distinct populations LA3111 (central) and LA4330 (southern highlands), with a conidial suspension of *C. fulvum* race 5. Upon visual inspection and microscopic evaluation, we found that plants of population LA3111 show similar phenotypes as our resistant control plant MM-Cf-9, a recombinant inbred line of *S. lycopersicum cv*. MoneyMaker with introgressed *Cf-9*, thus being resistant to *C. fulvum* race 5 as this race does produce Avr9. By contrast, LA4330 shows a phenotype similar to our susceptible control MM-Cf-5, a *Cf-5* introgression line of *S. lycopersicum* of which resistance is circumvented due to a loss of the Avr5 gene by race 5 (Mesarich et al., 2014) (Figure 1a-b).

**Figure 1:**
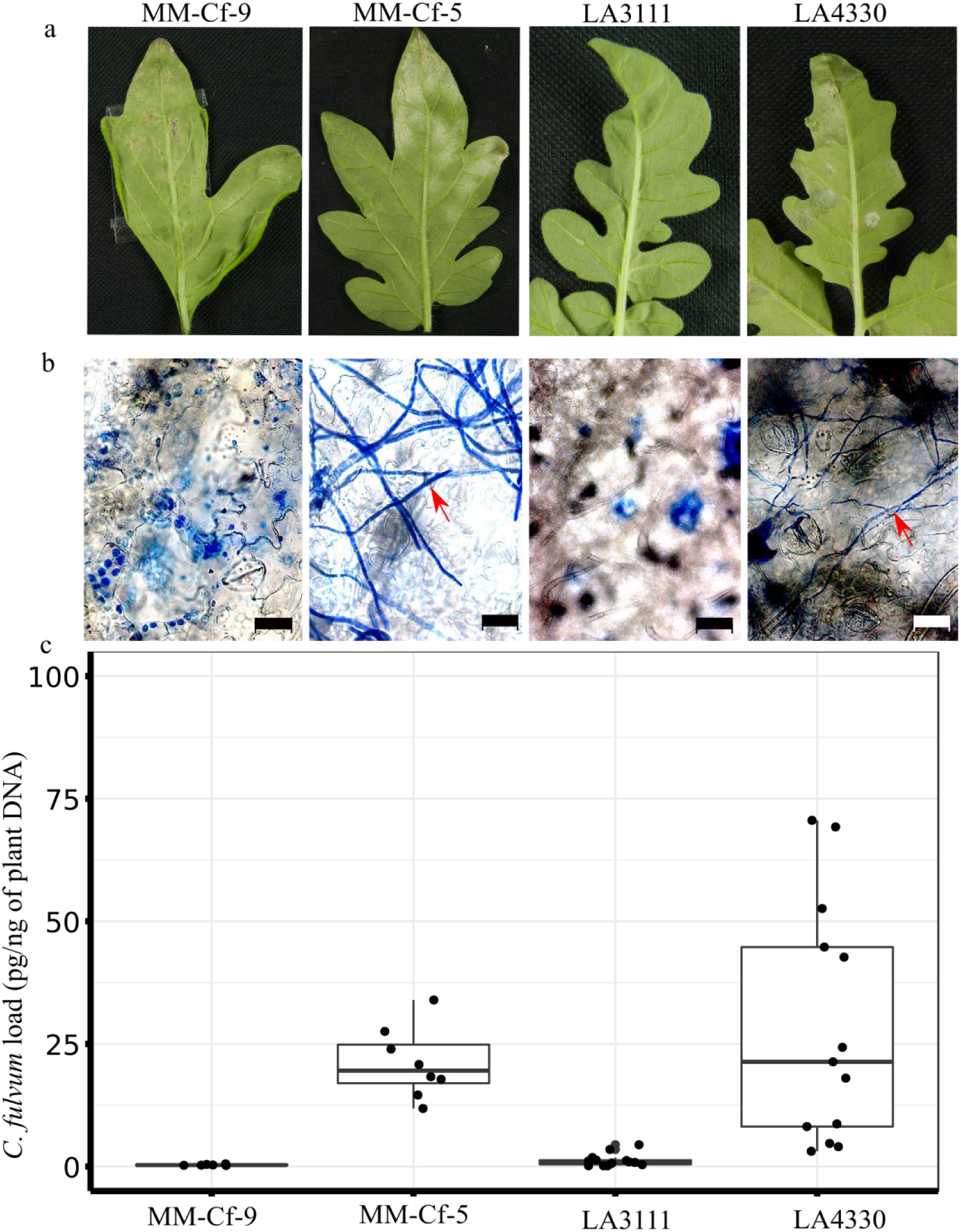
Tomato leaves inoculated with 2×10^4^ conidia/ml. a) Left to right: inoculated leaves harvested at 14 dpi of MM-Cf-9, MM-Cf-5 and *S. chilense* population LA3111 and LA4330. b) Microscopic pictures of bleached leaves (scale bar = 20µm), after staining with ink (mycelium of pathogen indicated with red arrows). c) Quantification of *C. fulvum* DNA load in pg/ng of plant DNA after 14 dpi of inoculation in 3 plants per MM control and 5 plants per *S. chilense* population. All the plants were evaluated in three technical replicates and each data point shows pathogen load per technical replicate.

Next we performed quantification of *C. fulvum* DNA load and found significant differences in pathogen DNA contents between the tested plants (p < 0.001, ANOVA). Differences can be seen between the resistant and susceptible control plants (p = 0.05, TukeyHSD) and LA4330 showed a significantly higher (p < 0.001, TukeyHSD) presence of *C. fulvum* DNA than LA3111 (Figure 1c). For this we tested five individual plants from each *S. chilense* population and three plants per control. Taken together these findings show that *C. fulvum* is able to infect *S. chilense* plants and that the various populations behave differently when exposed to *C. fulvum.*

### Southern *S. chilense* populations do not recognize *C. fulvum* effectors

Visible HR upon effector infiltration is an efficient and reliable proxy to test for the resistance properties of tomato plants on a large scale. To test the geographical variation in *C. fulvum* race 5 resistance, we performed an infiltration assay with apoplastic fluid (AF) of infected susceptible tomato plants, which is sufficient to trigger an HR in plants carrying matching *Cf* genes. As for the inoculation a race 5 was used, the AF is anticipated to contain all secreted *C. fulvum* effectors, except for Avr5. We infiltrated the AF at two sites in the leaves of 155 individuals, representing 15 different populations (8-17 individuals for each population). Interestingly, we did not observe any HR-associated recognition of an effector present in the AF by any of the 6 southern populations that we tested, e.g. those belonging to the southern highlands and southern coastal groups as described by Böndel (2015) and Stam *et al*. (2019b) (Figure 2). Populations from the northern and central regions do show recognition of at least one of the effectors present in the AF, as HR development was taking place. Yet, we did not observe this recognition to take place in all tested plants of the population. Rather, we observed differences in recognition capacities within the population, with some plants being able to respond with an HR, and some not. Populations from central regions showed recognition of components present in the AF, ranging from 10-100% of the plants tested in the different populations (Table S1). Two northern populations tested showed AF mix recognition for 75% and 80% of the plants tested (Table S1). All of our infiltration experiments were performed at least three times, in leaves from the same perennial plants and at the same location.

**Figure 2:**
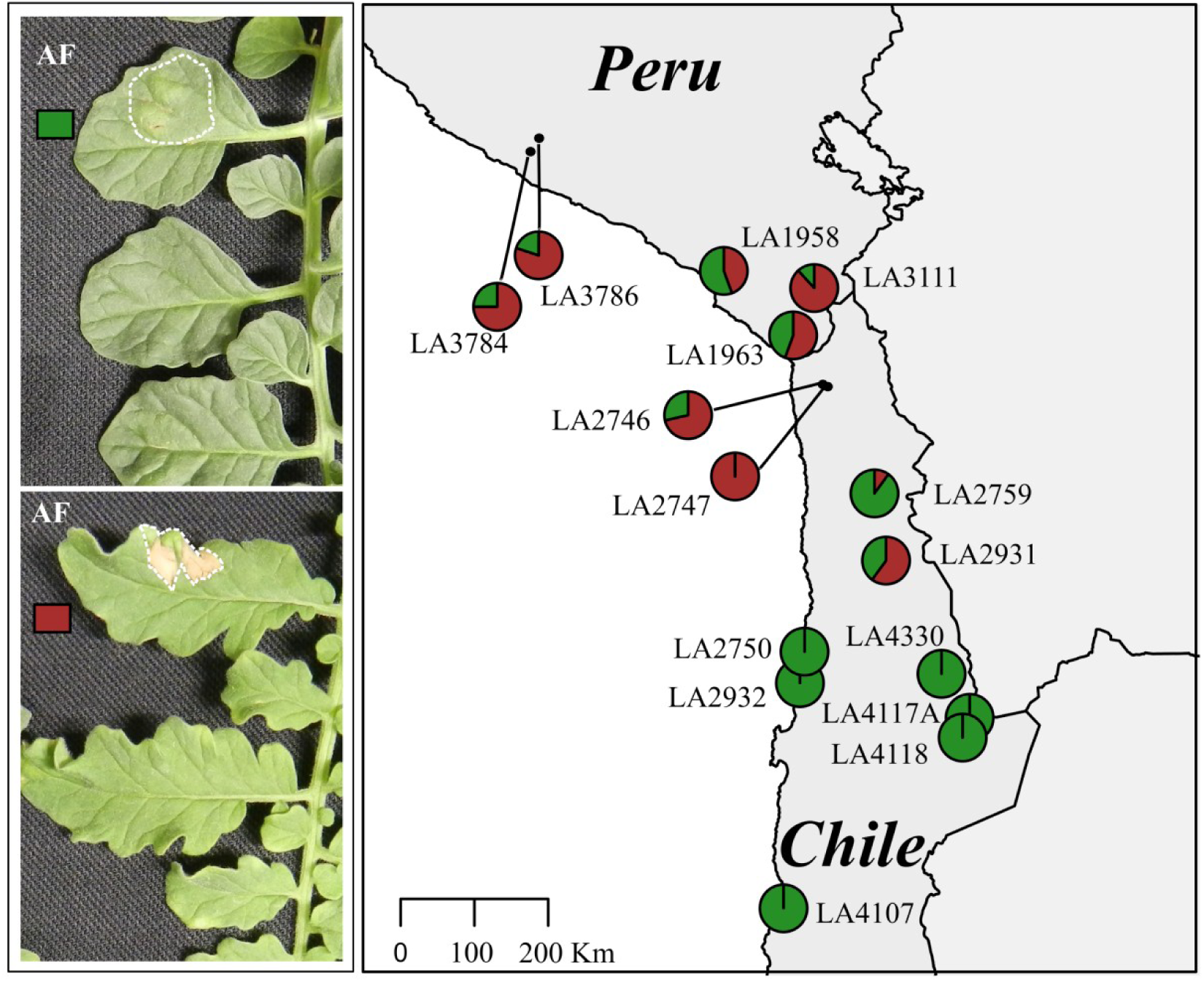
Infiltration of AF in populations of *S. chilense*. Leaf of an individual plant with no recognition of components present in the infiltrated AF (top left) and leaf of an individual plant recognizing at least one component present in the AF (bottom left). The infiltrated areas are indicated with white dotted lines. The map on the right shows the geographical distribution of AF component perception in *S. chilense*. The map shows the geographically distinct populations of *S. chilense* and their response to infiltration with AF Each pie chart indicates one population with 8 to 17 individuals. The brick red fraction represents plants that respond to *C. fulvum* AF, whereas green indicates the fraction of non-responding plants.

### Differential Avr9 and Avr4 recognition patterns are present in *S. chilense*

Recognition of Avr9 and Avr4 has been hypothesized to be an important conserved feature in wild tomato species in order to maintain *C. fulvum* resistance (Kruijt *et al*., 2005). To understand the role of the recognition of these Avrs in more detail, we performed additional infiltration assays, but now with the individual effectors. In order to test whether *S. chilense* plants are able to specifically recognize Avr9, we infiltrated the plants with a preparation enriched for Avr9. Populations from the central region showed a very large variation in the ability to recognize Avr9, with the capacity to recognize Avr9 ranging from 10-80% of the plants that were tested in individual populations (Table S1). The two northern populations that were tested showed Avr9 recognition for 20% and 37.5% of the plants tested (Figure 3).

**Figure 3:**
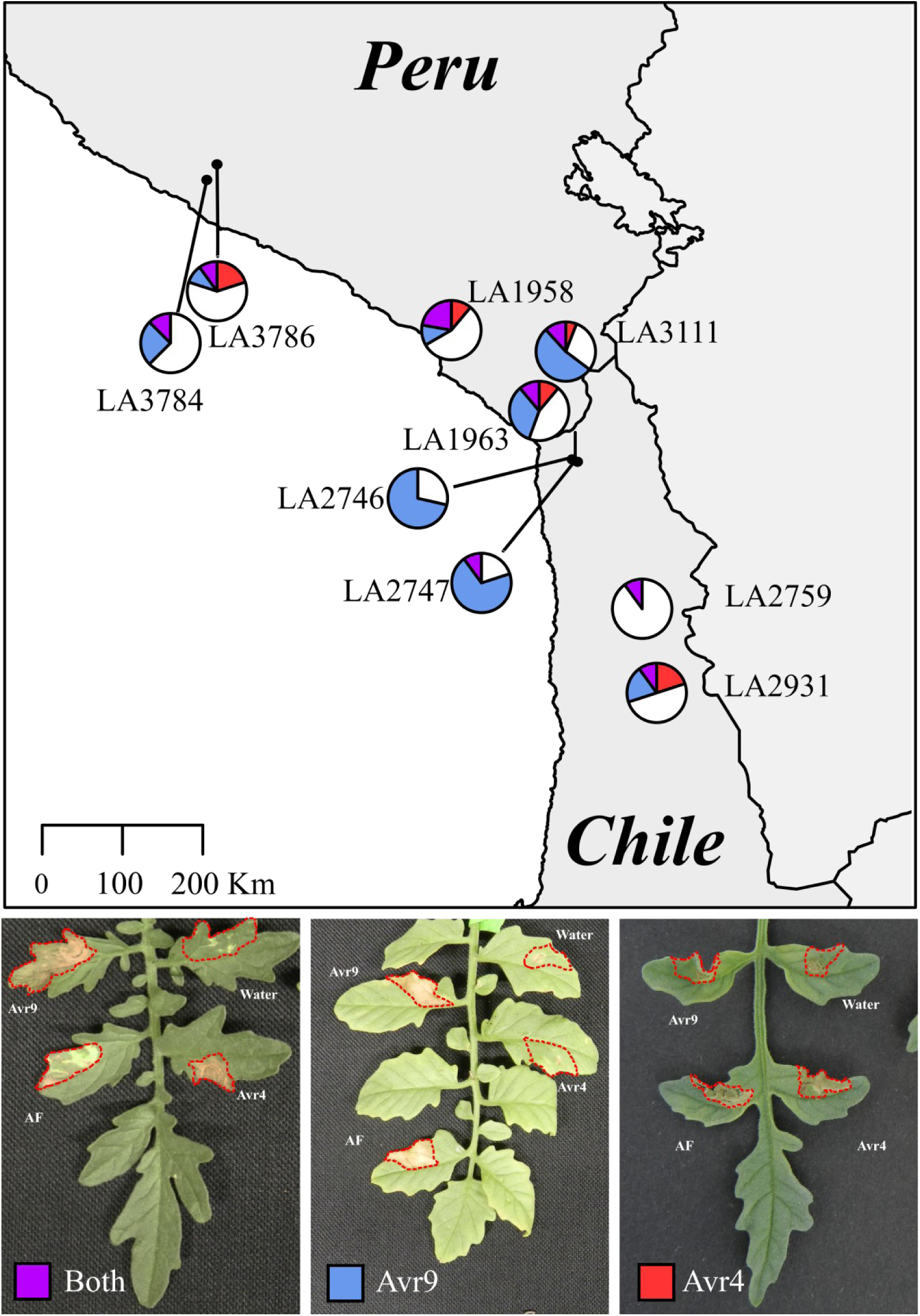
Geographical distribution of Avr9 and Avr4 perception in *S. chilense*. The map shows populations of *S. chilense* and their response to infiltration with Avr9 and Avr4 (purple when there is an HR-associated response to both), Avr9 (blue when there is an HR-associated response) and Avr4 (red when there is an HR-associated response). White sectors indicate plants showing no recognition of either Avr9 or Avr4. The infiltrated areas are indicated with red dotted lines. Each pie chart indicates one population with 8 to 17 individuals.

We also tested the ability of *S. chilense* to recognize Avr4 in a similar fashion, using *Pichia pastoris*-produced Avr4. In the two northern populations we observed that 12.5% and 30% of the plants showed the capacity to recognize Avr4, whereas in the central populations 0-33% of the plants showed recognition. Interestingly, contrary to previous reports that *Cf-4* and *Cf-9* are mutually exclusive, we also found plants that were able to recognize both Avr4 and Avr9 (Figure 2 and Table S1).

### Presence of canonical *Cf-9-, 9DC-* and *Cf-4*-specific region in different *S. chilense* populations does not correlate with their recognition properties

It has been shown that in *S. pimpinellifolium* either *Cf-9* or *9DC* are responsible for Avr9 recognition (Van der Hoorn *et al*., 2001). *Cf-9* or *9DC* gene sequences in *S. chilense* have not been reported to date. Putative full length *Cf-9* or *9DC* genes cannot be found in the currently available reference genome sequence, possibly due to misassemblies of the complex LRR regions (Stam *et al.*, 2019a). Thus, to investigate which gene is responsible for Avr9 recognition in *S. chilense* we looked into the presence of *Cf-9* and/or *9DC*, using gene-specific primer sequences that were used before to identify these genes (Van der Hoorn *et al.*, 2001).

We performed *Cf-9* gene-specific PCR amplification on genomic DNA isolated from nine *S. chilense* populations (9 plants per population), using the previously published primers CS5-CS1, amplifying a 379bp specific region of *Cf-9* (Van der Hoorn *et al.*, 2001). *Cf-9* and *Cf-4* introgression lines of *S. lycopersicum* served as positive and negative control, respectively. The *Cf-9*-specific sequence was amplified from all plants, although with different efficiency from all plants from the phenotyping assay. However, there was no association between presence or abundance of the amplicon and the response to Avr9 (Figure 4a).

**Figure 4:**
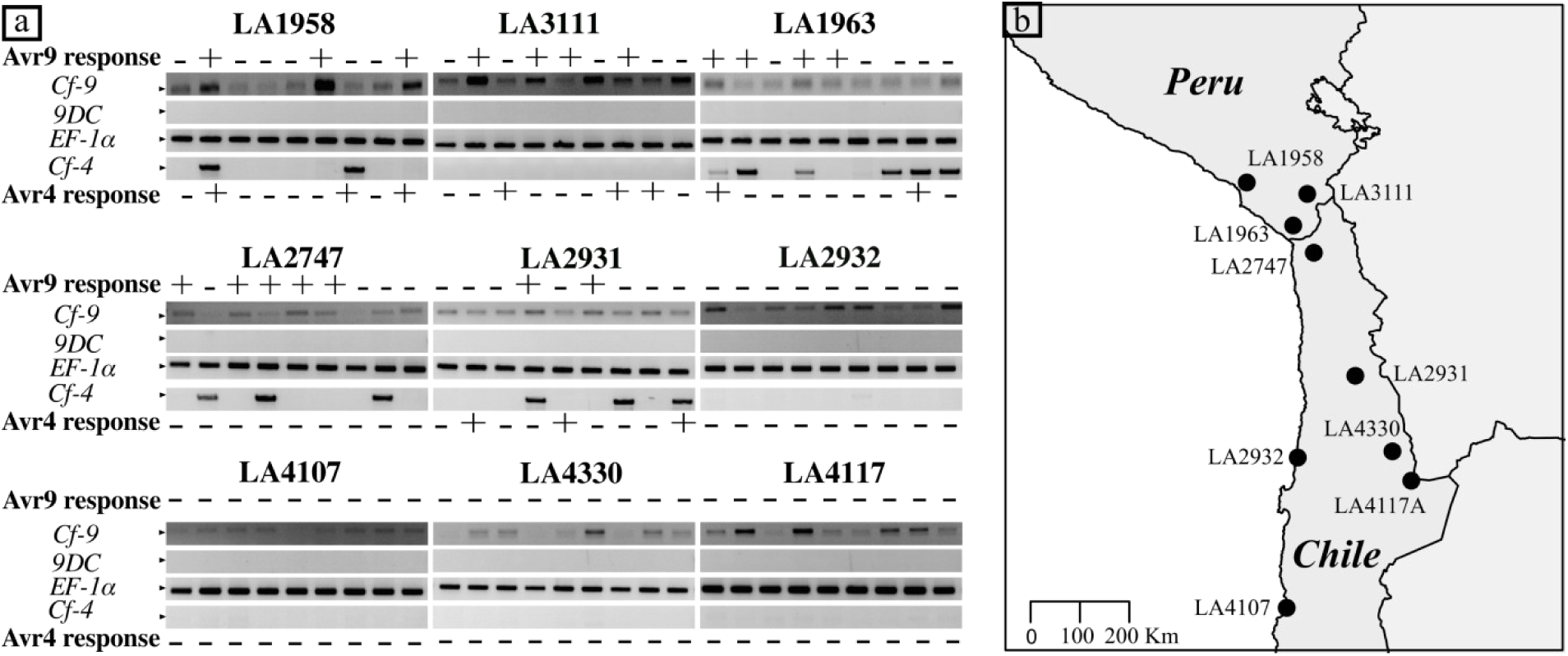
Amplification of the canonical *Cf-9, 9DC* and *Cf-4* region in geographically distinct populations of *S. chilense*. a) The CS5/CS1 primer pair was used to amplify a fragment of *Cf-9* (378 bp), the DS1/CS1 primer pair for *9DC* (507bp) and the PSK047/PSK050 primer pair for *Cf-4* (728bp). As a PCR control part of the coding region of *elongation factor 1 alpha (EF-1α)* was amplified using the primer pair RS003/RS004 (400 bp). MM-Cf-9 and MM-CF-4, and LP12 (*9DC*) were used as controls (Figure S2). + or – are indicative of Avr9 and Avr4 responsiveness upon infiltration of these effectors (Figure 2). b) Map shows the populations used in the analysis.

In a similar way, we tested the presence of *9DC* using the gene-specific primers DS1-CS1 (product size 507bp). LP12 and MM-Cf-9 served as positive and negative control, respectively. We found complete absence of the *9DC* canonical region in our populations (Figure 4a). Lastly, we also evaluated the presence of *Cf-4*, using newly designed primers (PSK047-PSK050) that fall over the intron that defines the difference between *Cf-4* and *Cf-9* (Figure S2) and amplify a 786bp product that is only present in *Cf-4*. We found that this *Cf-4*-specific region is present in a few plants belonging to the central population, suggesting that unlike in other *Solanum.* spp., *Cf-4* and *Cf-9* are not mutually exclusive in *S. chilense* (Figure 4). In addition, the presence of bands corresponding to the canonical *Cf-4* region in plants non-responsive to Avr4 and the absence in responders, suggests that other or new recombinant gene products with differences in their function exist in the different populations.

### *Cf-9* is expressed in a southern population

To test whether failed *Cf-9*(*-like*) gene expression is responsible for the complete loss of Avr9- triggered HR in the southern populations; we extracted RNA from 10 individuals from the LA4330 population. We used a primer pair PSK009-PSK010 that should allow us to amplify cDNA originating from *Cf-9* orthologs and performed semi quantitative RT-PCR on plants infiltrated with water and Avr9 (8 hours after infiltration), assuming that Avr9 recognition might upregulate *Cf-9*(*-like*) gene expression.

Without treatment, we found a transcript expressed in all tested *S. chilense* plants, as well as in our MM-Cf-9 control plant (Figure S4), indicating that steady state *Cf-9* gene expression is not affected in the southern *S. chilense* populations.

As expected, in the MM-Cf-9 plant we see a stronger band at 8 hours after treatment, indicative of the upregulation and positive feedback induced by *Cf-9*, a trait that is not visible in the southern *S. chilense* population (Figure S4).

### Loss of resistance might be due to mutations in *Cf*-coreceptors the southern populations

Seeing that steady state expression levels of *Cf-9* are not affected in the southern populations and complete loss of resistance is unlikely to result from detrimental mutations on all individual *Cf* genes. We hypothesize that the general loss of *Cf* responses likely results from changes in *Cf* regulatory genes, rather than mutations in the individual *Cf* genes themselves. Several core regulators or co-receptors are known to regulate the Cf protein function.

To test this hypothesis, we examined genome sequence data that are available for three plants, which are from representative populations of the central, southern coastal and southern highlands region (LA3111, LA2932 and LA4330 respectively) (Stam *et al*., 2019a). We extracted and aligned the genomic sequences of the co-receptors *SERK3a* (also known as *BRI1-ASSOCIATED KINASE 1, BAK1*) and *SOBIR1*, the adaptor *ACIK1*, required for regulation of the complex, as well as another regulatory co-receptor of the *BAK1*-containing complex, *BIR2* and its homolog *BIR2b* (Table S2). In all cases, the extracted sequence data show polymorphisms between the reference genome of *S. lycopersicum* Heinz1706 and the Avr4 and Avr9 responsive plant from LA3111. In-frame indels were found for *BIR2b* between its sequence in *S. lycopersicum* and those in the three *S. chilense* populations, yet, the three *S. chilense* populations show similar, complete sequences. Several unique polymorphisms exist in the southern populations, e.g. occurring in the plants from LA2932 and LA4330, but not in LA3111 from the central region or in the *S. lycopersicum* reference genome sequence, resulting in up to 19 non-synonymous amino acid changes for *SERK3a* in the plant from LA4330 (Table 1). In *ACIK1*, both the LA2932 and the LA4330 plant have unique indels not present in the *S. lycopersicum* reference genome sequence and in the LA3111 plant (Table 1). These results suggest that there are various possible amino acid changes these receptors, co-receptors and adaptors that could be responsible for the observed loss of resistance.

**Table 1:** Number of non-synonymous SNPs or indels observed in the open reading frame of genes encoding co-receptors or adaptors of *Cf* genes in the two southern populations LA2932 and LA4330.

| Gene Name | SNPs in<br>LA2932 | SNPs in<br>LA4330 | SNPs in both<br>LA2932 and LA4330 |
| --- | --- | --- | --- |
| *SOBIR1* | 4 | 1 | 0 |
| *SERK3a* | 4 | 19 | 1 |
| *ACIK1* | 1 | 3 | 2 |
| *BIR2* | 2 | 3 | 0 |
| *BIR2b* | 2 | 1 | 1 |

## Discussion

Natural plant populations are hypothesized to maintain a certain durable resistance against naturally co-occurring pathogens, yet little is known about the dynamics and the underlying genomic diversity in these populations throughout a species range, which result in the maintenance or loss of durable resistances.

*C. fulvum* is likely a natural pathogen of wild tomato species and known functional *Cf* resistance genes have been isolated from several wild tomato species. Yet, the physiological interaction between *C. fulvum* and wild tomato has not been documented. We show that *C. fulvum* race 5 is able to infect *S. chilense.* The formed intercellular hyphae resemble those observed in cultivated tomato, meaning that also infection in nature will lead to severe reduction in photosynthetic potential and thus will lead to loss of host fitness. We observed, both by the naked eye, as well as using microscopy and staining, that after inoculation with *C. fulvum* plants from a *S. chilense* population from the southern edge of the species range (LA4330) show higher susceptibility when compared to plants from a population from the central part of the range (LA3111). These findings were confirmed via quantification of fungal DNA. Thus, we show a compatible interaction between *C. fulvum* and wild tomato species, but also clear differences in resistance between the host populations.

The interaction between *C. fulvum* and tomato has been proven to be governed by gene-for-gene interactions, in which secreted Avrs from *C. fulvum* are recognized by corresponding RLP product of *Cf* gene from tomato (Joosten & de Wit, 1999; Stergiopoulos & de Wit, 2007). We tested recognition of such Avrs in different populations of *S. chilense* from different geographical locations with diverse climatic conditions, to understand whether the observed resistance and susceptibility within *S. chilense* populations to *C. fulvum* follows a specific geographical pattern. We phenotyped fifteen populations of *S. chilense* covering the whole species range by infiltrating an apoplastic extract potentially containing all Avrs except for Avr5, which is able to elicit immune responses that result in an HR. In northern and central populations, around 70% of the plants recognized at least one Avr present in this extract. Populations from the southern highlands and the southern coastal genotype groups showed no Avr recognition. These results point at two important conclusions. First of all, Avr recognition in the northern and central regions is not as conserved as previously hypothesized and secondly, Avr recognition appears to have been completely lost at the southern edge of the species range.

Previous reports suggest that Avr9 and Avr4 are the main factors in recognition of *C. fulvum* and this recognition is conserved throughout all wild tomato species (Kruijt *et al*., 2005). We observe Avr9 recognition to be present in 29% of the tested plants, whereas 11% of the plants recognized Avr4. These findings are somewhat contrasting with earlier results. Kruijt *et al*. (2005) tested only a small number of plants per population and did not report on the actual amounts of responding plants. In line with our results, they showed Avr4 recognition in 16 out of 20 tested populations and comparison of the accession numbers and linkage to their geographical origin, confirmed that only four of the populations tested by Kruijt *et al.* belonged to the southern genotype groups. Surprisingly, Avr9 recognition was not observed in all 20 accessions of *S. chilense* tested by Kruijt *et al*. (2005), although similar Avr9 have been used. These differences in results could be explained by the fact that all previous reports used young seedlings to allow quick screening of the plants. In our assays we use fully mature (over 1-year-old) adult plants, and we performed repeated infiltrations of the same plants. In addition, we observed differences in the strength of the HR upon Avr9 infiltration in young and fully developed mature leaves (Figure S3). Our finding therefore poses the first evidence of Avr9 recognition in *S. chilense* and suggests that detailed testing of other species under different conditions might yield novel interesting results. In about 6% of the tested plants we now show dual recognition of both Avr4 and Avr9, which has not been shown earlier.

It has been shown that Avr9 can be recognized by two *Cf-9* homologs, referred to as *Cf-9*/Hcr9- 9C and the recombinant *9DC*. For both of them, allelic variants with only a few nonsynonymous mutations are known. We evaluated the presence of known canonical domains that define *Cf-9* and *9DC* and found that the tested plants in all our populations do have the *Cf-9* domain but are lacking the *9DC* domain. The sole presence of the *Cf-9* domain is an interesting contrast with the findings of Van der Hoorn *et al*. (2001), who showed the presence of *9DC* to be predominant in another wild tomato species, *S. pimpinellifolium*, and a complete loss of *Cf-9* in southern populations of that spoecies. Note that the southernmost *S. pimpinellifolium* populations are geographically relatively close to the most northernly *S. chilense* populations that were analyzed in our current study, yet they populate clearly different ecological niches (Peralta *et al.*, 2008).

Likewise, we also evaluated the presence of the *Cf-4* canonical region in our populations. The canonical *Cf-4* domain is present in some individuals, but this does not correlate with our phenotypic data on the development of an HR upon infiltration with the Avr4 protein. There can be various reasons for this observation. First, the annealing sites of the *Cf-4* primers used might carry crucial SNPs in *S. chilense*, resulting in failure of the PCR no detectable bands. Second, the targeted region might be missing in the gene coding for the Avr4-responsive Cf protein, which would not be surprising as similar mechanisms of recombination events, resulting in the generation of a new gene of which the encoded protein has retained its recognition specificity, have been reported earlier for other *Hcr9s*, albeit on a phylogenetic rather than population scale (Van der Hoorn *et al*., 2001). Moreover, a study on *Cf-2* identified 26 different homologs in *S. pimpinellifolium* populations and revealed possible presence/absence variation of *Cf-2* among individuals (Caicedo & Schaal, 2004). Presence of canonical regions of *Cf-4* in the individual with *Cf-9* canonical region confirms the dual presence of previous though allelic gene which can be possibly due to heterozygosity at the locus.

Recombination and gene conversion have been shown to play a major role in gene family evolution for RLP family as well as other resistance gene families (Paniske *et al.*, 1997; Mondragon-Palomino & Gaut, 2005; Mondragon-Palomino *et al.*, 2017). Recently, it has been reported that such gene conversions, or micro recombinations, of NLR genes can also be observed between accessions of another wild *Solanum* sp., leading to an alternative mechanism to maintain different functional alleles (Witek *et al.*, 2020). We conclude that also the *Cf* gene family is likely not conserved *sensu stricto*, and hypothesize that a large number of possibly functional alleles are formed and maintained through intragenic micro recombinations, not just on a phylogenetic scale, but also between or even within populations of the same species.

The loss of Avr recognition in southern populations could be explained by the possibility that these plants recognize different Avrs which are absent in *C. fulvum* race 5. Yet, our findings show that northern and central populations of *S. chilense* do possess resistance against *C. fulvum* race 5. Our apoplastic extract contains at least 70 secreted effectors, which are all potential Avrs (Mesarich *et al.*, 2017). The chance of losing recognition for all of these effectors by mutations in the matching receptors is highly unlikely, thus hinting at general differences in the regulation of Avr recognition. Semi quantitative RT PCR experiments showed that *Cf-9*-like genes appear to be expressed in both Avr9-recognizing and non-recognizing plants. Single point mutations in genes encoding essential co-receptors, like SOBIR1 or BAK1 have been shown to lead to loss in ability to induce HR when expressed in a heterologous system (Bi *et al*., 2016) or *Arabidopsis thaliana* (Albert *et al*., 2019). Thus accumulation of deleterious mutations in such coreceptors or contrarily, accumulation of beneficial mutations related to, for example drought adaptation, that generate a trade of in resistance, are possible. BAK1/SERK3a is a co-regulator, in not only defense responses, but the receptor-like kinase also plays a role in general cell regulatory processes and is drought-responsive (Schwessinger *et al*., 2011). Genomic data for non-responding plants from the southern populations revealed several nonsynonymous mutations, as well as indels in multiple RLP/Cf co-receptor-encoding genes, when comparing them to either the sequence from the LA3111 reference plant, or the *S. lycopersicum* reference Heinz 1706, thus suggesting that changes in downstream regulation mechanisms could have caused the loss of Avr recognition in the south. Follow-up studies should determine which of the many altered genes are causal. Since the co-receptors have multiple functions apart from Cf protein signaling, such experiments might simultaneously shed light on how the different components of the signaling network interact.

Loss of resistance over time at the population level is rather poorly understood, with some theories possibly explaining this mechanism (Koskella, 2018). For instance, evolutionary loss of resistance in certain populations might happen as a result of genetic drift, e.g. after a severe bottle neck. However, if these populations would encounter any pathogen pressure afterwards, this would be detrimental. Loss of resistance can be the outcome of random processes that are triggered to take place due to loss of selection pressure, but this loss can also be the result of accelerated evolution due to the, much debated, assumed fitness costs of carrying obsolete resistance genes (Tian *et al*., 2003; Sheldon & Verhulst, 1996). The role of distribution of *C. fulvum* in shaping the evolutionary distribution of *Cf-2* homologs has been proposed by Caicedo (2008).

Giraud *et al.* (2017) point out that parasites need to develop a local adaptation to their environment beyond their host. Loss of resistance, as observed in our system, strongly suggests that a mechanism of ecological feedback is taking place, where the ecology of the population becomes a driving force to lose resistance or to maintain it. Therefore, the absence of the *C. fulvum* pathogen in the southern locations, due to extremely dry climatic conditions, which are not suitable for infection by *C. fulvum*, would be a plausible explanation. Interestingly, higher susceptibility to other pathogens also requiring relatively high humidity for successful infection, like *Phytophthora infestans* and *Alternaria* sp., was already described for southern populations of *S. chilense* (Stam *et al*., 2017). Seeing that some co-receptors potentially play a role in abiotic stress responses, a third possibility would be that the loss of resistance comes as a tradeoff for environmental adaptation. Yet, this is likely still intrinsically coupled to a decreased pathogen pressure.

In conclusion, we show that *Hcr9* locus in *S. chilense* is much more complex than was thought before (Parniske *et al*., 1997; Van der Hoorn *et al*., 2001; Kruijt *et al*., 2004; 2005). This might in part be due to the stronger niche differentiation, or the larger heterozygosity of *S. chilense* specifically (Moyle, 2008). However, it is also likely that there is a lot of undiscovered diversity in the *Cf* gene family present in other *Solanum* species. Furthermore, we provide an example of the loss of resistance in a wild tomato species at the edge of its geographical distribution, possibly explained by changes in the underlying immune receptor complexes.

Overall this study provides a step forward in terms of placing major gene-mediated molecular defense mechanisms in an ecological context.

## Acknowledgements

We like to thank Liza Keitel and Lina Muñoz for help with the experiments, Sabine Zuber, Bärbel Breulmann and Anneliese Keil for maintaining the *S. chilense* populations and members of SBF924 for fruitful discussions and useful feedback.

## Author contributions

Conceptualization: RS, RH, PSK and MJ; Investigation: PSK, MSS, GZ and DS; Contribution of materials: MJ; Funding acquisition: RS; Writing: PSK and RS. All authors reviewed and approved the manuscript.

