## supplemental data for "Population studies of the wild tomato species *Solanum chilense* reveal geographically structured major gene-mediated pathogen resistance"

**Supplemental Information: Kahlon et al 2020**

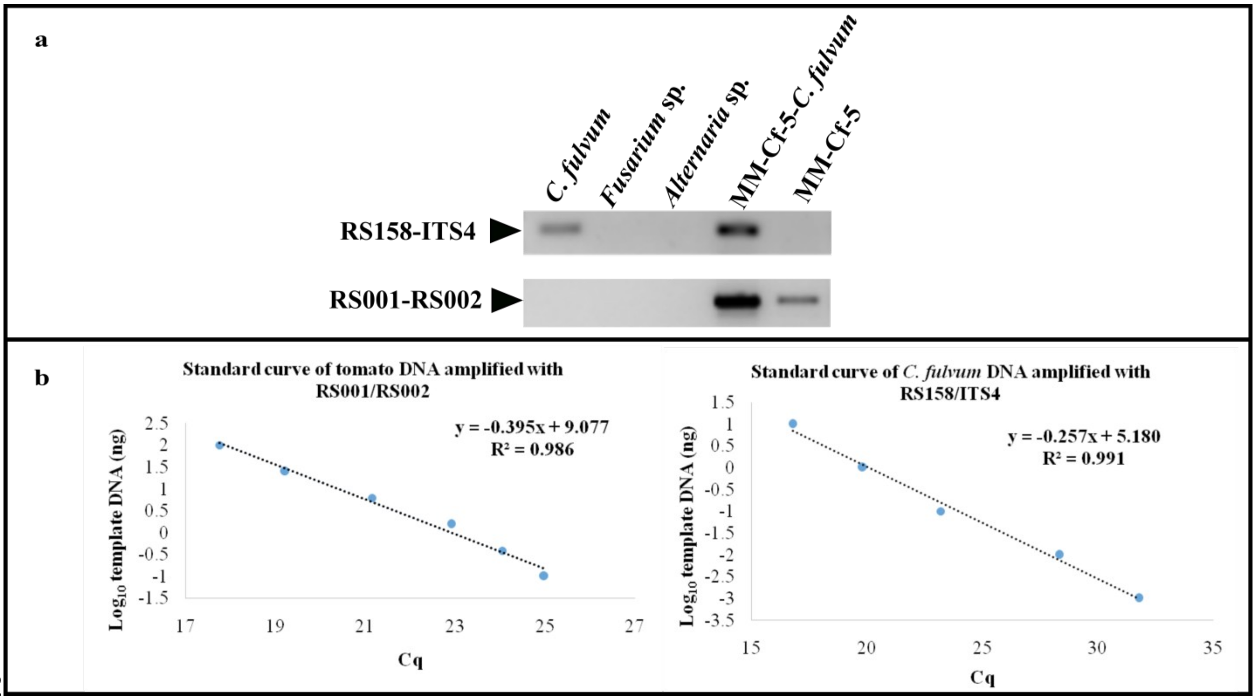


Figure S1: a) Primer pair specificity of RS158-ITS4 used to amplify the *C. fulvum* ITS region and specificity of primer pair RS001-RS002, used to amplify the alpha-tubulin fragment., b) regression line obtained upon plotting of the log10 values against the ITS and alpha-tubulin Cq values of serial-diluted *C. fulvum*(right) and tomato (left) DNA. The plotted Cq value represents the average of two technical replicates.

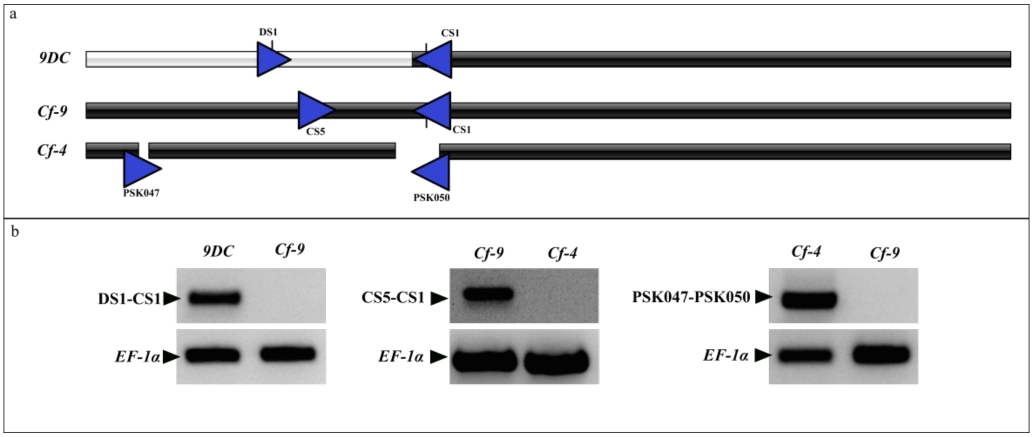


Figure S2: a) *9DC*, *Cf-9*and *Cf-4* alignments and primer annealing sites (blue triangles) (adapted from van der Hoorn *et al.*, 2001), b) PCR amplification products run on agarose gel, obtained by PCR with the different primer pairs indicated in (a), on genomic DNA obtained from LP12 (carrying *9DC*) and the introgression lines *Cf-9* and *Cf-4* used to evaluate the specificity of the primer pairs. The lower panel shows the PCR controls amplifying *elongation factor 1 alpha (EF-1α)* (RS003/RS004).

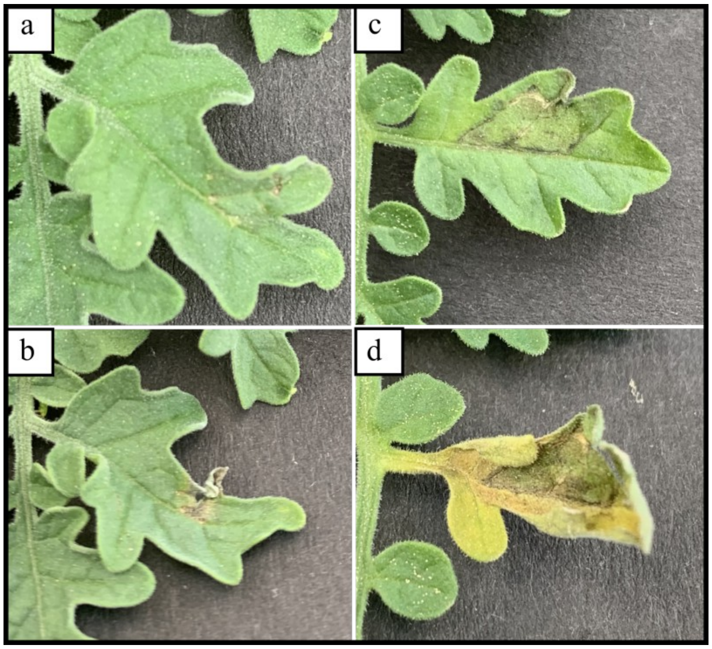


Figure S3. . The response to Avr9 is dependent on the age of leafthat is infiltrated. a) and b); young, first fully developed leaf infiltrated with Avr9, showing no HR at four days post infiltration (a) and HR appears when observed at day 7 (b); c) and d);an older leaf of the same plant shows HR within four days post infiltration with Avr9, in the same experiment.

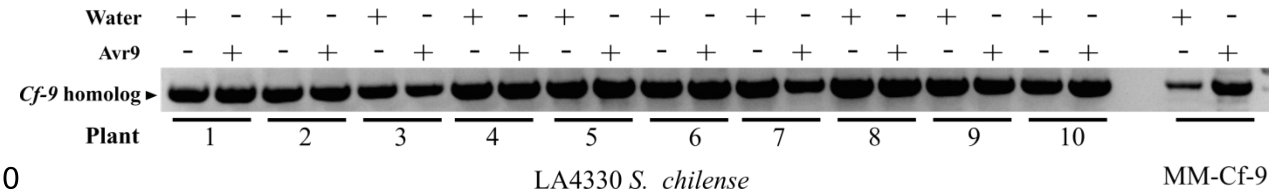


Figure S4: Semi quantitative RT-PCR of cDNA originating from a *Cf-9* homolog, amplified with primer pair PSK009-PSK010 (Table S3) of 10 individuals from population LA4330 and the *Cf-9* introgression line as a positive control, eight hours post infiltration with water or Avr9.

Table S1: AF, Avr4 and Avr9 recognition in different populations of *S. chilense*.

| Population | Region | Number of plants showing HR upon infiltration of | | | | Total plants tested |
|---|---|---|---|---|---|---|
| | | AF | Avr9 only | Avr4 only | Both Avr9 and Avr4 | |
| LA3784 | North | 6 | 2 | 0 | 1 | 8 |
| LA3786 | North | 8 | 1 | 2 | 1 | 10 |
| LA1958 | Center | 4 | 1 | 1 | 2 | 9 |
| LA1963 | Center | 5 | 3 | 1 | 1 | 9 |
| LA2746 | Center | 10 | 10 | 0 | 0 | 14 |
| LA2747 | Center | 10 | 7 | 0 | 1 | 10 |
| LA2759 | Center | 1 | 0 | 0 | 1 | 10 |
| LA2931 | Center | 6 | 2 | 2 | 1 | 10 |
| LA3111 | Center | 15 | 9 | 1 | 2 | 17 |
| LA2750 | S. coast | 0 | - | - | - | 9 |
| LA2932 | S. coast | 0 | - | - | - | 10 |
| LA4107 | S. coast | 0 | - | - | - | 10 |
| LA4117A | S. mountains | 0 | - | - | - | 9 |
| LA4118 | S. mountains | 0 | - | - | - | 12 |
| LA4330 | S. mountains | 0 | - | - | - | 8 |

Table S2: Known co-receptors and adaptors of Cf proteins and their annotation in *S. lycopersicum* and *S. chilense*.

| Gene Name | *S. lycopersicum* | *S. chilense* |
|---|---|---|
| *SOBIR1* | *Solyc06g071810* | *SOLCI002627000* |
| *SERK3a* | *Solyc10g047140* | *SOLCI001000400* |
| *ACIK1* | *Solyc07g041940* | *SOLCI000527600* |
| *BIR2* | *Solyc02g087460* | *SOLCI003763100* |
| *BIR2b* | *Solyc02g067560* | *SOLCI005193700* |

Table S3: Sequences of the primers used for qPCR and PCR in this study.

| Sr.No | Name | Sequences ('5-NNNN-3') |
|---|---|---|
| qPCR | | |
| 1. | RS001_Tubulin_Forward | GCCTACCATGAGCAGCTTTC |
| 2. | RS002_Tubulin_Reverse | CAATGCGTGAGAAGACCTCA |
| 3. | RS158-ITS-Forward | GTCTCCGGCTGAGCAGTT |
| 4. | ITS4-ITS-Reverse | TCCTCCGCTTATTGATATGC |
| PCR | | |
| 5. | RS003-EF1A_-Forward | GTCCCCATCTCTGGTTTTGA |
| 6. | RS004-EF1A-Reverse | GGGTCATCTTTGGAGTTGGA |
| 7. | DS1-*9DC*-Forward | GAGAGCTCAACCTTTACGAA |
| 8. | CS5-*Cf-9*-Forward | TTTCCAACTTACAATCCCTTC |
| 9. | CS1-*Cf-9-9DC*-Reverse | GCCGTTCAAGTTGGGTGTT |
| 10. | PSK009-*Cf*-Forward | ATGGATTGTGTAAAACTTGTATTCCT |
| 11. | PSK010-*Cf*-Reverse | CTAATATCTTTTCTTGTGCTTTTTCA |
| 12. | PSK047-*Cf-4*-Forward | ACGACAGAAGAACTC |
| 13. | PSK050-*Cf-4*-Reverse | GATGGAATTGGTCCTT |
